# Visuo-tactile integration in texture perception: A replication and extension study

**DOI:** 10.1101/2022.04.01.486675

**Authors:** Karina Kangur, Martin Giesel, Julie M. Harris, Constanze Hesse

## Abstract

Employing a sensory conflict paradigm, previous research has found that vision and touch contribute, on average, equally to the visuo-tactile perception of surface texture. Our study aimed to, firstly, replicate the original findings using a comparable setup and stimulus set; secondly, examine whether equal modality contributions can also be observed on an individual basis (using a within-subject design); and thirdly, explore how visuo-tactile integration is affected by illumination angle (top vs. oblique). Participants explored a discrepant standard consisting of different abrasive papers by vision, touch, and using both modalities simultaneously, and subsequently had to find the closest visual, tactile, and visuo-tactile match from a set of matching stimuli. We replicated equal contribution from vision and touch across the whole sample in both illumination conditions. We also found considerable inter-individual variations in the modality contributions when the stimuli were illuminated from the top. Interestingly, this variation decreased under oblique illumination, with most participants showing an about-equal contribution from both modalities to the combined texture percept. These findings are consistent with the assumption that the perceived discrepancy between vision and touch was reduced under oblique illumination suggesting overall that visual and tactile information are only weighted equally within a certain range of experienced discrepancy. Outside this individual range, one of the modalities is weighted higher with no clear preference for either modality.

## Introduction

Texture perception is complex as it is multidimensional as well as multisensory [1]. Surface roughness is one important dimension of 3D texture and material perception that has been studied extensively (e.g., [2-4]). Perceived roughness is a composite texture property that depends on a number of different physical parameters such as the size, density, spacing, and jaggedness of the particles on surfaces [5, 6] which may be evaluated visually, haptically, and/or auditorily. Consequently, many studies have focussed on how sensory information is weighted and integrated across different modalities to produce a unified percept of 3D texture (for a review, see [7]). Here, we investigated the integration of visual and haptic information of surface properties with a focus on inter-individual variations in the way modalities are weighted.

Studies examining visual and haptic judgements of surface roughness consistently show that both measures are highly correlated (e.g., [2, 8]) and that judgements are similarly accurate when made by either vision or touch [5, 9, 10, 11]. Interestingly, in contrast to other visuo-haptic object properties (e.g., object size, 12]) as well as other modality combinations (e.g., audio-visual signals [13], or gustatory and olfactory stimuli, [14]), these studies found no compelling evidence for multimodal enhancement, that is, improved performance when textures were presented bimodally as compared to unimodally. In other words, bimodal performance is not found to be generally superior to unimodal performance (but see [15] for a different view). This finding has led to the suggestion that vision and touch may act as independent sources of roughness information [7, 10].

The finding of vision and touch being similarly accurate in providing information about roughness further means that it cannot be easily determined if the different sources of visual and haptic information are integrated, or if information from one modality is ignored in situations where congruent information is available to both senses simultaneously. One way to resolve this issue is to use a sensory discrepancy paradigm in which an artificial conflict is introduced between the information presented to different sensory modalities, seemingly originating from exploring the same physical event (e.g., [16, 17]; for a review, see [18]).

Lederman and Abbott [5] used such a discrepancy paradigm to study the relative contributions from vision and touch to texture perception when performing matches to abrasive papers either by vision, touch or using both modalities together. The results of their Experiment 1 showed that when abrasive papers of different roughness (i.e., different grit values) were explored by vision and touch, the combined texture matches were approximately the mean of the unimodal texture matches. Equal contributions from visual and tactile modalities have also been observed in subsequent studies using abrasive papers [11] and fabric samples [10]. These findings can be well described by a weighted averaging model ([1], see Method for details) where visual and tactile information are assigned approximately equal weights. Assuming similar precision for visual and haptic texture perception [5, 10, 11], the equal weighting of visual and tactile input in the discrepancy paradigm might also indicate statistically optimal integration as predicted by the maximum-likelihood integration model [12] in which modality weights are assigned in a way that minimises the variance in the combined percept.

However, despite the evidence for equal contributions from vision and touch, it has been shown that there can be great variability in modality weights between participants when making visual and haptic judgments about certain object properties, such as size, form, or texture (e.g., [10, 16, 19]), with some participants showing complete visual or haptic dominance.

Since Lederman and Abbott [5] employed a between-subject design where for each exploration and matching condition a different group of participants (N=10) provided only one single match/trial (i.e., nine experimental groups), they were unable to determine whether the equal weights for vision and touch veridically represented individual weightings or if they partly resulted from combining the data from visually and tactually dominant participants. It has been suggested that individual differences in modality weights may also be linked to the precision with which individuals can make visual and tactile judgements (for reviews see [7, 18]) as well as their experience and personal sensory preferences (e.g., [20-22]). However, overall, studies investigating the relationship between sensory acuity, experience and modality biases are sparse and their findings are inconclusive.

Hence, the main objective of our study was to replicate and extend Experiment 1 of Lederman and Abbott’s study and to establish whether equal modality weightings in a discrepancy matching task are also consistently observed for individual participants. Consequently, we applied a within-subject design which allowed us to determine visual and tactile weights for each participant separately.

Furthermore, Lederman and Abbott [5] illuminated their stimuli from the top. As there is evidence that roughness perception is not independent of illumination angle and direction [23-24], we were also interested in exploring the role illumination may play in visuo-tactile integration when using discrepant roughness stimuli. More specifically, it seems that lowering the illumination angle results in the surfaces being perceived as rougher. To our knowledge, the only study that systematically varied illumination angles while measuring visual and tactile judgements of surface roughness found that the sensitivity and efficiency of visual judgments improved under oblique illumination conditions [21]. At the same time, visual judgements became more similar, in terms of accuracy, to tactile ones. Brown speculated that participants’ ability to discriminate visual roughness improved under oblique illumination due to increased grain-shadowing. This speculation is in line with Ho et al. [23] who found that patterns of shading and cast shadows may be among the visual cues that participants use to make roughness judgements.

To explore the potential effect of illumination on texture perception in a discrepancy paradigm, we employed two different illumination conditions. In Experiment 1, the discrepant standard and matching stimuli were illuminated directly from the top. In the following, we will refer to this as *top illumination* (i.e., illumination perpendicular to the surface of the stimuli). We assumed that the illumination of Experiment 1 was similar to the illumination condition used by Lederman and Abbott and thus represents a close replication of their study. In Experiment 2, we lowered the illumination to a grazing angle (*oblique illumination*) which allowed us to test if visual roughness perception indeed varies with illumination angle, and whether and how this affects the integration (and respective weights) of visual and tactile information. Finally, we also assessed participants’ tactile acuity [25] as well as their personal preferences for haptic information processing (Need for Touch Scale, [26]), in order to explore if any potential individual differences in those measures correlate with modality preferences (as measured by modality weights) in a discrepancy matching task.

## Materials and methods

### Participants

Twelve participants took part in Experiment 1 (6 males, M_age_ = 29.2 years, SD_age_ = 5.3 years, age range: 24-40), and ten of those also completed Experiment 2 (5 males, M_age_ = 29.2 years, SD_age_ = 5.5 years, age range: 24-40). The sample size was chosen to match that of Lederman and Abbott [5] where each participant performed one single trial in one of the exploration and matching conditions (10 participants (i.e., trials) per condition and 90 participants (i.e., trials) in total). As we employed a within-subject design, we aimed to recruit at least ten participants to match the number of trials per condition from Lederman and Abbott’s study.

Near visual acuity was measured using the SLOAN letter chart (40 cm distance, SKU: 52185, Good-Lite: 756400) and was at least 20/20 for all participants. The tactile acuity thresholds were smaller than 3 mm for all participants as determined by the JVP domes (Stoelting Co., Wood Dale, IL, USA).

The study has been approved by the local ethics committee of the School of Psychology at the University of Aberdeen (PEC/4360/2019/10) and was performed in accordance with the ethical guidelines of the British Psychological Society. All participants provided written informed consent and were reimbursed £30 for completion of Experiments 1 and 2.

Note that the methods and data analysis procedures of this study have been pre-registered on the OSF prior to starting data collection, and are publicly stored with the data associated with this project: https://osf.io/nzb78/ (DOI 10.17605/OSF.IO/NZB78)

### Discrepancy matching task

#### Stimuli and setup

Our stimuli were identical to those used by Lederman and Abbott [5] consisting of nine different abrasive papers (7.5 cm x 12.5 cm) with the following grit values: 40, 50, 60, 80, 100, 120, 150, 180, 220. Grit values refer to the number of openings per square inch in the sieve used to apply the particles to the papers [27], with lower grit values referring to rougher, and higher grit values to finer, abrasive papers. To eliminate colour cues (and following Lederman & Abbott’s procedure), all abrasive papers were painted with a single coat of black semi-gloss enamel paint. Abrasive papers were cut in half (3.75 cm x 6.25 cm) to create a visual and a tactile stimulus set. There were nine matching stimuli for which the visual and tactile grit values were identical. The discrepant stimulus (we will call this the *discrepant standard*) was a combination of a 150 grit paper for visual exploration and a 60 grit paper for tactile exploration.

The setup consisted of a black rectangular box (H: 60 cm, W: 73 cm, D: 30 cm, Fig 1A) placed on a table with two openings: the upper one (W: 15 cm, H: 10 cm) for the visual and the lower one (W: 15 cm, H: 5 cm) for the tactile exploration of the stimuli (comparable setup to Lederman & Abbott [5]). A rotating table (IKEA SNUDDA, d = 39 cm) was placed inside the box in line with the tactile exploration opening, and the nine matching stimuli and the discrepant standard were placed on it. The tactile stimuli were arranged in a circle around the perimeter of the rotating table, and the visual stimuli were placed in an adjacent inner circle (Fig 1B). The matching stimuli were ordered counterclockwise from low to high grit values. The construction of the box prevented the participants from seeing the tactile and/or touching the visual stimuli while also ensuring that only one visual and/or tactile stimulus could be explored at a time.

**Fig 1.**
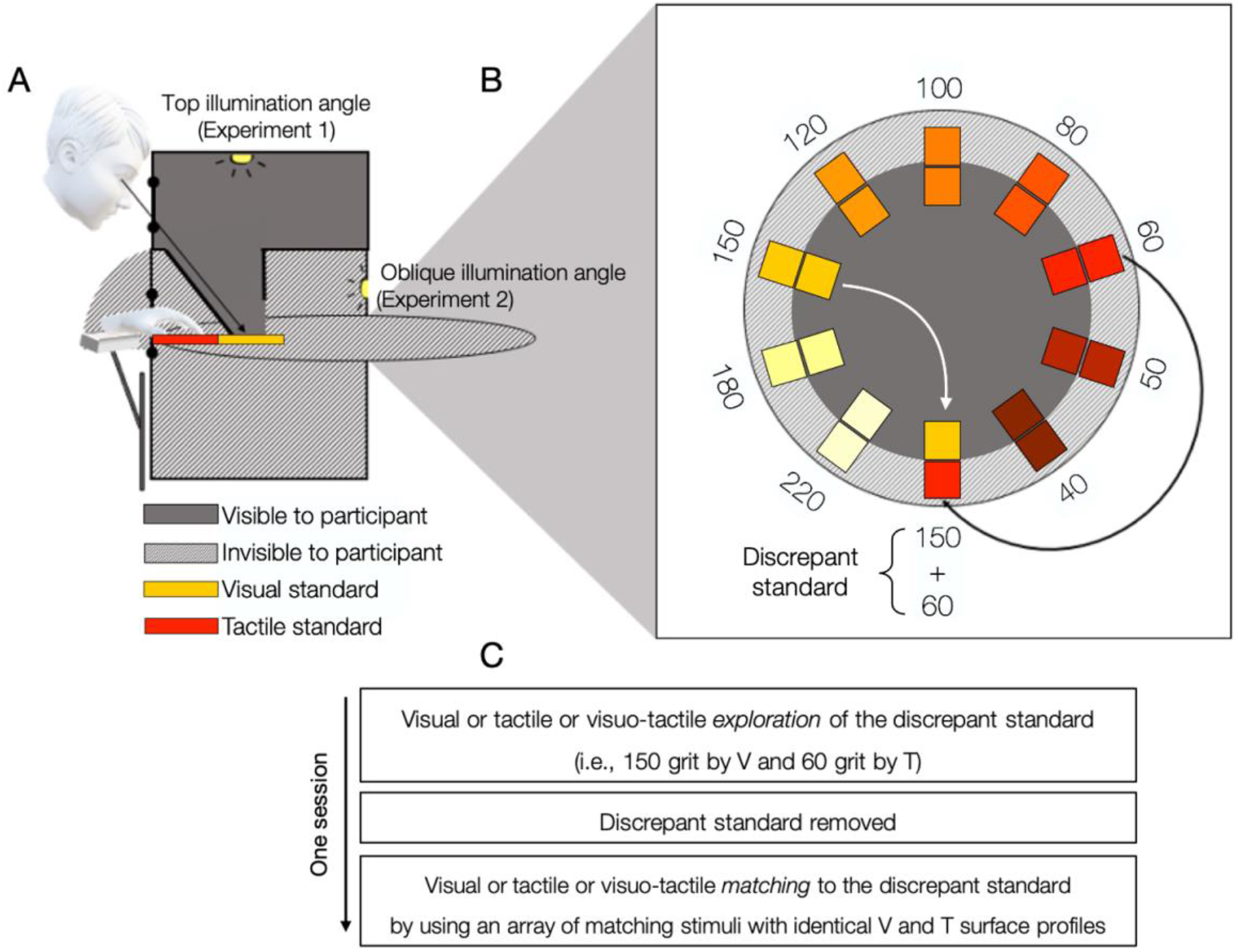
Setup of the discrepancy matching task. (A) An example of the trial where the discrepant standard was explored on the rotating table visually and tactually (visuo-tactile exploration). Please note that while this image depicts both illumination angles, they were tested in separate experiments. The lights were not directly visible to the participants. (B) Stimuli and their ordered arrangement on the rotating table. Numbers indicate the grit values associated with the abrasive papers. The stimuli arranged around the perimeter were for tactile exploration while those around the adjacent inner circle were for visual exploration. The discrepant standard comprised the visual (150 grit) and the tactile standard (60 grit). The colours are for illustrative purposes only. (C) Experimental procedure for all sessions of the discrepancy matching task.

The visual stimuli were illuminated by a strip of three LED lights (colour temperature: 6000K) spaced at 1.5 cm intervals. In Experiment 1 (top illumination), the visual stimuli were illuminated from the top, whereas in Experiment 2 (oblique illumination), they were illuminated from an oblique angle (30 degrees). The viewing distance from the eye to the stimulus was 24 cm.

### Procedure

The matching task involved three exploration and three matching conditions: visual (V), tactile (T), and visuo-tactile (VT). This resulted in nine combinations of exploration and matching conditions: V-V, V-T, V-VT, T-V, T-T, T-VT, VT-V, VT-T, and VT-VT. Note, when we use these abbreviations, the letter(s) before the hyphen refer(s) to the exploration condition and the letter(s) after the hyphen to the matching condition.

Participants were first asked to explore the discrepant standard uni- or bimodally and then to pick a match to the standard from the array of matching stimuli uni- or bimodally. In the unimodal exploration conditions (i.e., starting with ‘V-’ and ‘T-’), participants explored the discrepant standard either by vision (always 150 grit) or by touch (always 60 grit) alone. In the bimodal exploration condition (i.e., starting with ‘VT-’), they explored the discrepant standard (always visually 150 grit and tactually 60 grit) using both modalities. In these bimodal trials, we ensured a simultaneous sensory presentation by lifting a visual cover the moment participants’ fingers made contact with the tactile standard.

Once finished with the exploration, the visual opening was covered and/or the participants removed their hand from the tactile opening. The experimenter then informed the participants whether matching would be done visually, tactually, or by using both senses together. Both the participant and the experimenter were unaware of the matching modality until after the participant had finished the exploration of the standard. Participants were asked to pick the matching stimulus that corresponded to the standard that they had just explored. During matching, only one stimulus was visible and/or felt at a time, and the participants were aware that the stimuli were arranged in the order of grit values. The exploration started randomly either from the roughest (i.e., 40 grit) or the finest (i.e., 220 grit) matching stimulus (following the procedure of Lederman & Abbott [5]). The experimenter moved the rotating table as per the verbal instructions from the participants (e.g., “next”, “go back”) until a match was chosen, thus concluding the matching trial (i.e., session). Please note that at no point was the term ‘roughness’ mentioned in the instructions of the experimenter, participants were simply asked to choose the closest match out of the available textures. Participants received no feedback about their performance at any point during the experiment.

The order of presentation of exploration and matching conditions was randomised. Each participant completed each of the nine exploration-matching combinations in different sessions, each consisting of one experimental trial (Fig 1C). To avoid possible learning effects, the sessions were completed on separate days. In the first session, participants’ visual and tactile acuity was assessed, they completed the haptic preference questionnaire (i.e., NFT Scale), and performed one matching trial. This session took just under one hour to complete. Consecutive matching sessions lasted no longer than five minutes, but no time restrictions were imposed during the exploration or matching.

Setup and procedure were identical in Experiments 1 and 2. Experiments only differed in the angle from which the stimuli were illuminated. Participants completed the first nine sessions in Experiment 1, followed by nine sessions with oblique illumination angle in Experiment 2. Since we assumed that the illumination in Experiment 1 closely resembles the illumination originally employed by Lederman and Abbott [5], the rationale for performing illumination conditions in separate (and subsequent) experiments was to separate the replication of Lederman and Abbott’s study from the more exploratory investigation of the effects of oblique illumination in this paradigm (see pre-registration).

### Deviations from pre-registered protocol

In the pre-registration (https://osf.io/nzb78), we suggested excluding participants who verbally commented on a perceived discrepancy in the visuo-tactile standard. Surprisingly, most of our participants commented on a potential sensory discrepancy at different points during Experiments 1 and 2. This was unexpected as Lederman and Abbott [5] stated that none of their participants reported having noticed a sensory discrepancy in a follow-up questioning despite using the same discrepant standard. Importantly, the detection of a sensory discrepancy in our experiments was neither due to accidental tactile exposure to the visual stimulus (as the experimenter watched participants’ hand movements during the exploration phase) nor to a lack of naivety about the purpose of the study (which none of our participants guessed correctly). We suspect that the prolonged exposure may have given more opportunities for the participants to detect and comment on a discrepancy. Since previous studies indicated that awareness of a sensory conflict has little or no effect on reported perception in discrepancy paradigms ([28]; for a review, see [18]), we decided against excluding participants based on these comments. Limitations that this could possibly impose on our results are discussed in more detail in the “Discussion” section.

### Data analysis

We manually recorded the numerical grit values of our participants’ matches. As outlined in the pre-registration document, the individual matches in Experiments 1 and 2 were analysed using a 3 (exploration modality: V, T, and VT) x 3 (matching modality: V, T, and VT) repeated-measures ANOVA. In line with Lederman and Abbott [5], significant main effects of exploration and matching modalities were followed up with the examination of the differences between the uni- and bimodal matches using paired-samples t-tests.

Some of the statistical comparisons we chose were not anticipated in the pre-registration, and thus not planned. For these non-planned (and labelled as such) comparisons, Bonferroni-Dunn corrections were applied. For all analyses, a significance level of α = 0.05 was used. Means are presented with ±1 SEM (between subjects).

Following Lederman and Abbott [5], we calculated the modality weights by averaging grit values over the three matching conditions. Equations 1 and 2 show how the weights for the visual (1) and tactile (2) modalities were computed (also see [5], page 906):

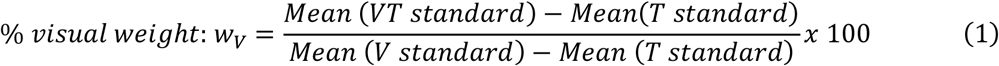

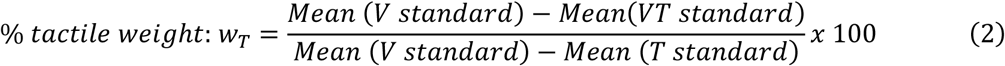

T refers to the mean grit values of all matches (visual, tactile and visuo-tactile) made following the exploration of the tactile standard, V to the mean of all matches made to the visual standard, and VT to the mean of all matches made to the discrepant visuo-tactile standard.

These equations were derived from assuming a weighted averaging model of visuo-tactile integration with the weights adding up to 1 (Equation 3; also see [1]).

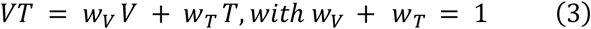

Using the within-subjects design allowed us to also calculate the individual visual and tactile weights: individual modality weights were based on the average of three matching trials per participant and condition. We used paired-samples t-tests to determine whether the weights of the two modalities significantly deviated from equal modality weighting.

Finally, to test if illumination affected matching performance, we conducted a 3 (exploration modality: V, T, VT) x 2 (illumination: top vs oblique) repeated-measures ANOVA. This analysis included the data from the ten participants who completed both experiments. We also determined the effect of illumination on the modality weights using a paired-samples t-test. Analyses were performed using JASP, R (RStudio IDE) and SPSS.

### Tactile acuity and haptic preference measures

Prior to the discrepancy matching task, we also assessed participants’ tactile acuity in a standardised grating orientation task (JVP Domes [25]), as well as their preferences for extracting and using haptic information using a standardised questionnaire (Need for Touch (NFT) Scale [26]). The tactile acuity task required participants to discriminate between vertical and horizontal orientations of differently spaced grating patterns of a small plastic dome (grating spaces of 0.35, 0.5, 0.75, 1.0, 1.25, 1.5, 2.0, 3.0 mm) applied to the index finger pad of participants’ dominant hand (2-AFC task). Each dome was administered 10 times (5 in each orientation) in random order resulting in a total of 80 trials. Tactile spatial acuity was determined as the 75% threshold of correct responses and provides an estimate of participants’ cutaneous spatial resolution. The NFT Scale consists of 12 questionnaire items that assess two dimensions of haptic experience (‘autotelic’ and ‘instrumental’). It requires participants to indicate their agreement on a 7-point Likert scale (‘Strongly disagree’ to ‘Strongly agree’). For both NFT scores and tactile acuity thresholds, we determined their correlation with the tactile weights in the discrepancy matching task.

## Results

### Experiment 1: Top illumination: Replication of Lederman & Abbott (1981)

We were interested in the differences in the matching grit values between the unimodal (i.e., visual and tactile only) and the discrepant bimodal (i.e., combined visuo-tactile) exploration conditions and whether those varied with matching modality. Based on the findings of Lederman and Abbott [5], we expected that the matches chosen after the exploration of the discrepant standard (i.e., VT-T, VT-V and VT-VT) would be roughly halfway between the matches made after unimodal tactile (60 grit) and unimodal visual (150 grit) exploration, i.e., around 105 grit.

Figure 2 shows the mean matches for the three exploration and matching modalities (2A) as well as the mean matches averaged across matching modalities (2B, i.e., the average for each of the three lines in 2A; note 2C and 2D show results for Experiment 2). As predicted, the average matches in the discrepant bimodal exploration condition were in-between the matches made after visual or tactile unimodal exploration for all three matching modalities. As expected, matches made after unimodal visual or tactile exploration were close to the grit values explored in those conditions (i.e., 60 grit tactile and 150 grit visual).

**Fig 2.**
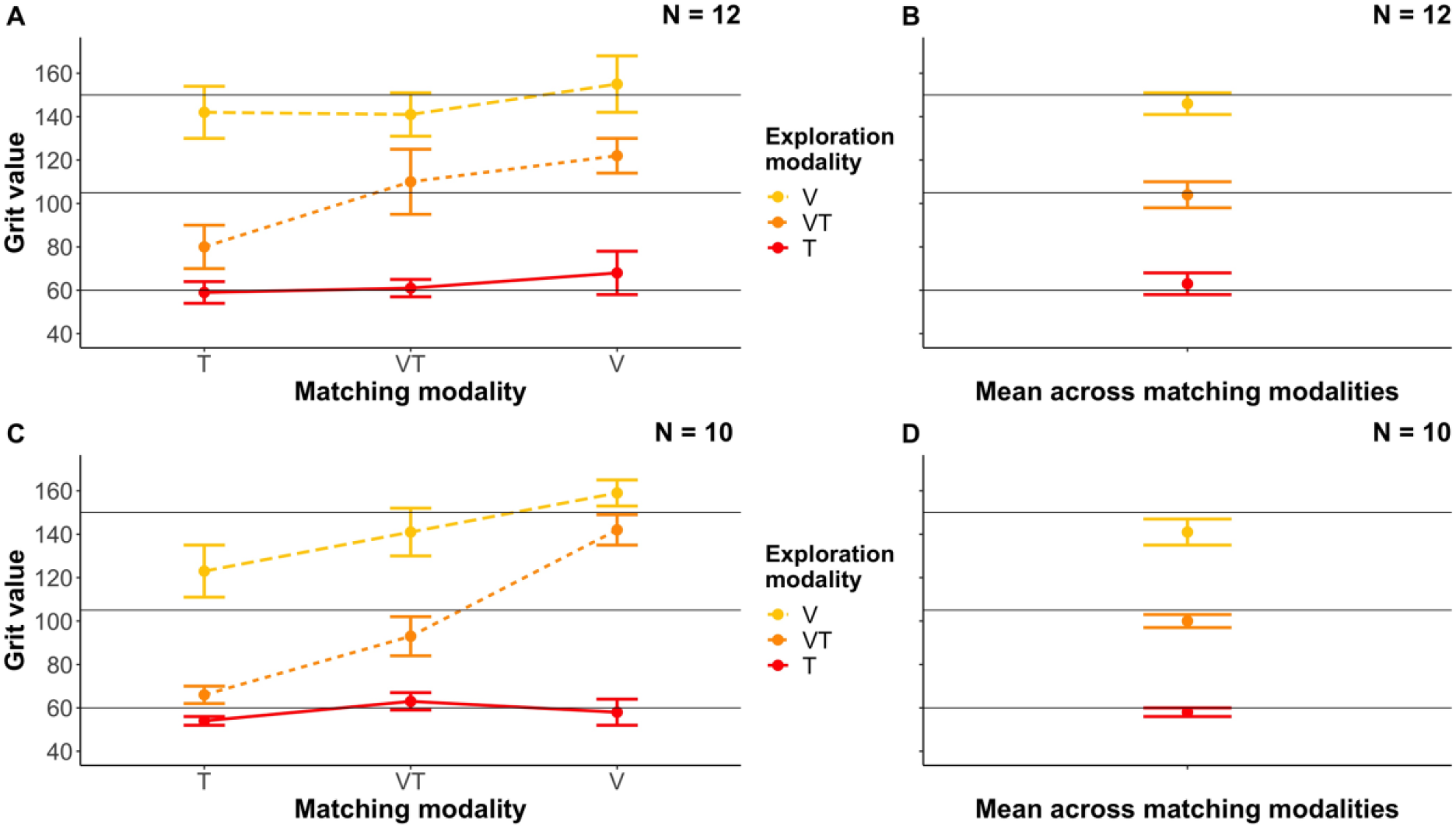
Mean matches in the top and oblique illumination experiments. Panels A and B refer to the top illumination (Experiment 1), C and D to the oblique illumination (Experiment 2) experiments. (A, C) Mean matches as a function of matching and exploration modalities. (B, D) Mean matches for the three exploration modalities averaged over the three matching modalities. Horizontal lines at 60 and 150 represent the grit values associated with the standards. Please note that the horizontal line at 105 grit represents the average of the visual (150 grit) and tactile (60 grit) standards. Error bars denote ±1SEM (between subjects). Figs A and C directly correspond to Fig 2 in Lederman & Abbott [5].

To test those observations statistically, we conducted a 3 (exploration modality: V, VT, T) x 3 (matching modality: V, VT, T) repeated-measures ANOVA on the individual matches. As predicted, our analysis revealed a main effect of exploration modality (*F*(2,22) = 55.25, *p* < .001, *η*_*p*_^*2*^= .834). Planned pairwise comparisons confirmed that the tactile standard was perceived as significantly rougher (63 ±5 grit) than the visuo-tactile standard (104 ±6 grit; *t*(11) = −5.00, *p* < .001), and the visuo-tactile standard as significantly rougher than the visual standard (146 ±5 grit; *t*(11) = 4.36, *p* = .001, Fig 2B). These results were expected as different standards were examined in the tactile (60 grit) and visual (150 grit) exploration conditions and are also in line with those reported by Lederman and Abbott [5]. One-sample t-tests further confirmed that there were no statistically significant deviations from the expected values for the tactile standard (60 grit; *t*(11) = 0.61, *p* = .61) and the visual standard (150 grit; *t*(11) = −0.83, *p* = .42). Furthermore, the matching values for the discrepant visuo-tactile standard did not differ from the average of the two unimodal standards (105 grit; *t*(11) = −0.13, *p* = .90). Contrary to Lederman and Abbott, the repeated-measures ANOVA revealed no significant main effect of matching modality, *F*(2,22) = 2.86, *p* = .079, *η*_*p*_^*2*^ = .206. The interaction between exploration modality and matching modality was also not significant, *F*(4,44) = 1.02, *p* = .41, *η*_*p*_^*2*^ = .085.

To calculate the relative modality weights in the discrepant condition, we first averaged the matches over the factor matching modality (i.e., three trials per participant and matching modality, resulting in one matching value per exploration modality). Then, weights were computed using the weighted averaging model (see Equations 1 and 2) suggested by Lederman and Abbott [5]. Figure 3 shows the averaged (3A) and individual weights (3C; note that 3B and 3C also show data for Experiment 2, see below). Averaged across all our participants, we found a mean visual influence of 52.4%, and thus a tactile influence of 47.6% (±12.5%, Fig 3A). Note that visual and tactile weights always add up to 100% using this model. These weights were very close to those reported by Lederman and Abbott who found a visual influence of 49.3% and a tactile influence of 50.7%. A paired-samples t-test showed that the difference between the visual and tactile weights was not significant, *t*(11) = 0.19, *p* = .85, further confirming an equal contribution from both modalities.

**Fig 3.**
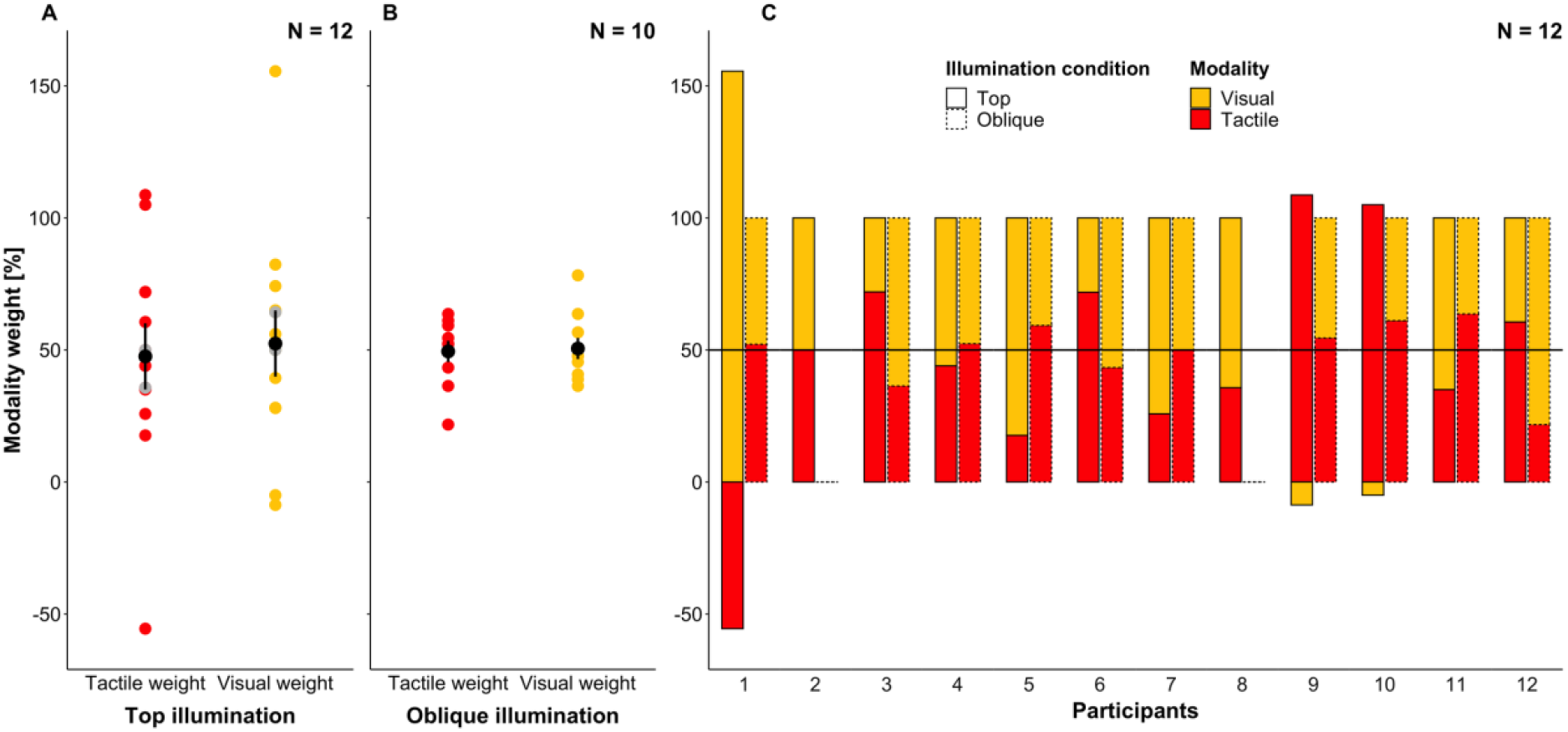
Tactile and visual modality weights in the two experiments. Panel A refers to top illumination (Experiment 1), panel B to oblique illumination (Experiment 2), and panel C to both experiments. Visual weights are *1 - tactile weights* and are plotted for illustrative purposes only. (A, B) Black dots denote the mean, and error bars denote ±1SEM (between subjects). Grey dots in A highlight the two participants who completed the top but not the oblique illumination experiment (participants 2 and 8). (C) The two experiments are denoted with different bar line types (top: solid line; oblique: dotted line).

Further, we aimed to determine if equal modality weights also occur on an individual level. As can be seen from Fig 3A (and solid bars in 3C), we observed clear individual differences in modality weights. It seems that while the two modalities are weighted about equally for some participants, others showed pronounced modality preferences. For three participants, we found weights that were outside the 0-100% range: two participants, 9 and 10, showed nearly complete tactile dominance, while participant 1 showed a visual weight well above 100%, and therefore a negative tactile weight (see Fig 3C). Weights outside the 0-100% range indicate that matches (i.e., grit values) for the discrepant visuo-tactile standard were outside of the range of matches that the participant selected for the unimodal visual and tactile standards. For example, Participant 1 chose an average match of 57 grit after exploring the tactile standard (60 grit), an average match of 117 grit after exploring the visual standard (150 grit) but an average match of 150 grit after exploring the discrepant visuo-tactile standard (60 grit tactually and 150 grit visually). This may potentially happen if the participant experienced a strong discrepancy between the visual and haptic stimuli and, therefore, assumed that the two sensory inputs emanated from different physical events ([29, 18], see discussion for more information).

Overall, our results replicate Lederman and Abbott’s [5] original findings confirming an approximately equal weighting of vision and touch in texture perception across participants. In addition, our findings reveal some inter-individual variations in the weighting of visual and tactile information. The weights found for some participants suggest that they may have experienced a discrepancy too large for visuo-tactile integration to take place.

### Experiment 2: Oblique illumination

In Experiment 2, we aimed to explore the effects of oblique illumination on texture/roughness perception and visuo-tactile integration. We expected that an oblique illumination angle would result in the visual standard and matching stimuli being perceived as overall rougher (see [23]). An increase in perceived roughness of the visual standard would result in a reduction of the perceived discrepancy between the visual and the tactile standard which, in turn, may affect the weights given to this information in the discrepant exploration condition.

Compared to the illumination from the top in Experiment 1 (i.e., top illumination), the illumination angle was lowered in Experiment 2 (i.e., oblique illumination). Otherwise, the experiment and data analysis remained identical to Experiment 1 (Fig 1).

Figure 2C shows the mean matching responses for the three exploration and matching modalities for Experiment 2. As in Experiment 1 (Fig 2A), the selected matches (averaged across matching modality) were close to the grit values of the visual and tactile standards after unimodal exploration, and in between those values after bimodal discrepant exploration (Fig 2D). The 3 (exploration modality: V, T, VT) x 3 (matching modality: V, T, VT) repeated-measures ANOVA on the individual matches revealed, again, a main effect of exploration modality, *F*(2,18) = 121.83, *p* < .001, *η*_*p*_^*2*^= .931. Planned post-hoc comparisons show that the matches to the tactile standard (58 ±2 grit) were significantly rougher than matches to the visuo-tactile standard (100 ±3 grit; *t*(9) = −9.0, *p* < .001). Moreover, matches to the visuo-tactile standard were significantly rougher than the matches to the visual standard (141 ±6 grit; *t*(9) = 9.4, *p* < .001, Fig 2D). One-sample t-tests showed that there were no statistically significant deviations of the matches for the tactile (i.e., 60 grit; *t*(9) = 0.69, *p* = .51) and the visual standards (i.e., 150 grit; *t*(9) = −1.61, *p* = .14) when compared to the actual grit values explored. Again, the average visuo-tactile match was also not significantly different from the average of the two unimodal standards (i.e., 105 grit; *t*(9) = −1.65, *p* = .13).

Similarly to the original findings by Lederman and Abbott [5] but contrary to our findings from Experiment 1, we found a significant main effect of matching modality, *F*(2,18) = 29.05, *p* < .001, *η*_*p*_^*2*^ = .763. However, this effect cannot be meaningfully interpreted as there was also a significant interaction between exploration and matching modality, *F*(4,36) = 4.95, *p* = .003, *η*_*p*_^*2*^ = .355. In order to understand the differential effects of matching modality on matches made in the different exploration conditions, we conducted post-hoc repeated-measures ANOVAs with the factor matching modality (V, T, VT) separately for each of the three exploration conditions. These analyses revealed that neither for tactile exploration, *F*(2,18) = 1.22, *p* = .32, *η*_*p*_^*2*^ = .12, nor for visual exploration, *F*(2,18) = 3.07, *p* = .07, *η*_*p*_^*2*^ = .25, were the matches significantly affected by the matching modality. In contrast, for the bimodal exploration of the discrepant standard, matching modality had a significant effect on the selected matches, *F*(2,18) = 25.10, *p* < .001, *η*_*p*_^*2*^ = .74. In the bimodal exploration condition, the unimodal matches (VT-T and VT-V) were closer to the grit values of the unimodal standards, respectively (see [5] for a similar observation and comparison between Experiments 1 and 2 for a more detailed discussion).

Modality weights are shown in Fig 3B. Again, we found on average an equal weighting of visual (50.5%) and tactile information (49.5% ±4%), *t*(9) = .14, *p* = .90, thus again replicating Lederman and Abbott’s [5] findings for oblique illumination. Interestingly, the inter-individual variability in modality weights seemed substantially reduced as compared to Experiment 1 (compare Fig 3A and B), and the weights of all participants were within the 0-100% range. To analyse the effects of illumination angle in more detail, we compared the matches and modality weights in the two experiments.

### Comparison of Experiments 1 and 2: Effects of Illumination angle on visuo-tactile integration

We compared how the matches and modality weights differed when the stimuli were illuminated from the top (Experiment 1) and from an oblique angle (Experiment 2) for those participants that finished both experiments (N=10).

Based on the work by Ho et al. [23], we expected the change in illumination angle in Experiment 2 to result in systematic changes in the visual appearance of the visual standard and the visual matching stimuli. Specifically, we expected the visual standard to appear rougher and, therefore, likely to be matched to a lower grit value tactually compared to the corresponding match in Experiment 1 (V-T condition). Additionally, the tactile standard may be matched to a smoother matching stimulus visually (T-V condition) than in Experiment 1 because of the overall rougher visual appearance of the matching stimuli under oblique illumination. Descriptively, tactile matches to the visual standard (V-T condition) decreased slightly in Experiment 2 (123 ±12 grit) as compared to Experiment 1 (137 ±14 grit), but this change was not statistically significant, *t*(9) = 0.74, *p* = .48. There was also no evidence that participants selected larger grit values visually after exploring the tactile standard (T-V condition: 70 ±12 grit in Experiment 1 vs. 58 ±5 grit in Experiment 2; *t*(9) = 1.04, *p* = .33).

To test if illumination and exploration modality affected the selected matches, we conducted a 2 (illumination: top, oblique) x 3 (exploration modality: V, T, VT) repeated measures ANOVA averaged across matching modality. As expected, this analysis revealed a significant effect of exploration modality, *F*(1,9) = 104.1, *p* <.001, *η*_*p*_^*2*^ = .92, indicating that matches were roughest for exploration of the tactile standard (61 ±4 grit) and least rough for exploration of the visual standard (143 ±4 grit), with matches to the discrepant standard falling in between the two (102 ±5 grit). However, there was no main effect of illumination and no interaction between the two factors (both p > .20, *η*_*p*_^*2*^ <.17), suggesting that participants made similar matches in both illumination conditions.

As discussed above, the most pronounced changes between the matches in the two illumination conditions were observed for the unimodal matches in the bimodal (VT) exploration condition (Fig 2, conditions VT-T and VT-V). To further explore this observation statistically, we performed a 2 (illumination: top, oblique) x 3 (matching modality: V, T, VT) repeated-measures ANOVA (none pre-registered analysis) on the matches made in the discrepant exploration condition (i.e., VT-). While there was no main effect of illumination, *F*(1,9) = 0.20, *p* = .66, *η*_*p*_^*2*^ = .022, the main effect of matching modality was significant, *F*(2,18) = 8.47, *p* = .003, *η*_*p*_^*2*^ = .49, as was the illumination x matching modality interaction effect, *F*(2,18) = 6.46, *p* =. 008, *η*_*p*_^*2*^ = .42. This interaction effect seems to be driven by the fact that in the oblique illumination condition, the unimodal matches (i.e., VT-T and VT-V) were closer to the grit values of the visual standard when matching visually and closer to the tactile standard when matching tactually than in the top illumination condition (i.e., VT-T: matched 82 grit under top and 66 grit under oblique illumination; VT-V: matched 117 grit under top and 142 grit under oblique illumination). Note that the observation that the matches selected for the discrepant standard are biased toward the modality with which the match is made, mirrors the findings and observations of Lederman and Abbott ([5], see their Fig 2).

Importantly, the finding that after bimodal exploration, participants select visual and tactual matches that are close to the respective visual and tactile standards, seems consistent with the assumption that the discrepancy between the visual and tactile standards was reduced under oblique illumination (i.e., the visual standard appears rougher and thus closer to the tactile standard). To make this reasoning clearer: Once participants had explored the discrepant VT-standard bimodally (i.e., using vision and touch simultaneously), they were asked to either match the stimulus using vision (VT-V) or touch (VT-T) only. If they experienced little or no discrepancy during exploration, they would be expected to select a match close to the respective unimodal standards (i.e., 60 grit haptically and 150 grit visually). If asked to select the match bimodally (VT-VT) participants would still need to resolve a mismatch given that the discrepant standard they had explored was not available in the array of matching stimuli. It is possible that the change in illumination angle that resulted in the decrease of the discrepancy in the standard at the same time also increased the discrepancy in the matching stimuli. The finding that the VT-VT matches remained largely the same for the two illumination angles is consistent with this assumption. However, as discussed in the introduction, assuming that visual and tactile senses are similarly accurate, it is impossible to tell for a non-discrepant standard whether the information is integrated or whether one of the senses is ignored when making a judgement as both would result in the same match. Yet, as we find integration under top illumination, there seems no good reason to assume that sensory dominance should occur under oblique but not under top illumination, since sensory dominance is more likely when a discrepancy is increased rather than reduced (e.g., [12, 18]).

Finally, we wanted to assess if the illumination angle affects the weights given to the two modalities. As shown in Figs 3A and 3B, on average, about equal modality weights for vision and touch were observed in both illumination conditions, *t*(9) = 0.059, *p* = .95 (wV = 51.5 ±15.1% in Experiment 1; and wV = 50.5 ±4.0% in Experiment 2; both N=10). However, what is notable (Fig 3C), is the reduced inter-participant variability when stimuli were illuminated from an oblique angle (Experiment 2). While a number of participants seemed to show some modality preferences in Experiment 1, with three participants showing behaviour inconsistent with a weighted averaging model, this did not occur in Experiment 2. To test this observation statistically, we conducted a Levene’s Test for the equality of variances for paired samples [30-31] on the haptic weights of Experiments 1 and 2 (none pre-registered test). This test confirmed that variances in modality weights were smaller in Experiment 2, t(9) = 2.32, p = .02 (one-sided).

Finally, we were interested in exploring whether individual modality weights correlated under the two different illumination conditions (none pre-registered analysis). Interestingly, we observed no significant correlation between modality weights, *r*(10) = −.11, *p* = .75. The fact that participants’ modality weights were uncorrelated in the two illumination conditions may suggest that these weightings do not relate to specific individual preferences or sensitivities but may instead more strongly be influenced by task-related/environmental factors.

This observation is further supported by the finding that tactile weights neither correlated with the spatial tactile acuity thresholds (as determined by the JVP Domes), despite relatively large individual differences (range: 0.68 to 2.38 mm, mean: 1.39 mm ±0.13, both *p* > .74) nor with the participants’ preferences in engaging in haptic interactions as determined by the NFT Scale (all p > .22). However, given the relatively small sample size, the lack of correlations should be interpreted with caution (see also pre-registration).

## Discussion

The main aim of our study was to test whether the almost equal weighting of modality contributions – as seen in Lederman and Abbott [5] on a group level – also holds on an individual level. Lederman and Abbott employed a between-subjects design meaning that weights could not be calculated for each participant separately but were instead determined across the whole sample. Consequently, it remained unspecified if the equal contribution from the two modalities occurred consistently across participants or may have been, in part, due to averaging responses across participants that were strongly biased either towards the visual or tactile modality (see [5], page 911, for discussion of this issue). Using the same task, stimuli and a comparable setup, but employing a within-subjects design, allowed us to compute individual modality weights and thus determine inter-individual differences. We also tested different illumination conditions to explore whether and how illumination affects texture perception and modality weights in a discrepancy paradigm. In both experiments, we found, on average, equal weighting of visual and tactile modalities. Hence, we replicated Lederman and Abbott’s findings consistently across both top and oblique illumination conditions.

Our exploration of individual differences provided some interesting new insights into the processing of discrepant visuo-tactile information. Specifically, we found larger inter-individual variations in the relative weights with illumination from the top (Experiment 1) with some participants showing a stronger preference for tactile and others for visual information. For one participant, we observed weights clearly outside of the expected 0-100% range and for another two participants tactile weights close to 100%. In contrast, for oblique illumination (Experiment 2), inter-individual variability was reduced, and all weights fell within the expected range.

It has been argued that multisensory integration depends on the amount of discrepancy that is introduced between the different modalities. As the experienced discrepancy becomes too large, one of the input modalities may be discounted resulting in visual or tactile dominance [12, 18]. This may have occurred for some participants in Experiment 1 thereby explaining the overall larger inter-individual variability. Furthermore, a number of participants mentioned that they noticed a discrepancy between the visual and tactile standards in Experiment 1. Previous literature suggests that awareness of a discrepancy alone has little effect on participants’ biases as long as they still believe that the same object/stimulus is explored (e.g., [18, 28]). However, if participants no longer believe that their visual and tactile sensations originate from the same physical stimulus, no inter-sensory discrepancy is experienced, and thus no integration is expected to occur. In this case, it is difficult to predict on what basis participants select their matches if the task requires them to do so, but it may result in modality weights that are no longer within the expected range (i.e., 0-100%). This might have been the case for participant 1 whose visual weight exceeded the 100% mark (see Fig 3C).

There are two potential explanations for the reduced inter-individual variability in Experiment 2. Firstly, as all participants completed the top illumination condition first (in order to separate the replication of Lederman and Abbott’s [5] study from the exploration of the effect of illumination angle), one could speculate that reduced variability may simply be due to training effects. However, even though this explanation seems quite straightforward, we deem it relatively unlikely for a number of reasons. First, participants only performed a total of nine trials under top illumination and never received any feedback about their performance in this condition. Second, all trials/sessions were performed on separate days, with participants completing all 18 trials/sessions over the course of several weeks. Third, variations between stimuli were relatively subtle. Whatever participants may have learnt in Experiment 1 about the mapping between visual and tactile experience and the range of the matching stimuli, might not have been applicable in Experiment 2 because of the change in the visual appearance of the standard and the matching stimuli. This argument is further supported by the lack of a correlation between the modality weights in the two experiments, suggesting that discrepant information may have been processed differently in the two tasks (i.e., dominance vs. integration).

Hence, we think that the reduced inter-individual variability in performance under oblique illumination may be better explained by the assumption that the perceived roughness of the visual standard was increased, which in turn would reduce the perceived discrepancy of the visuo-tactile standard. In particular, we found that after bimodal exploration, the unimodal tactile matches were close to the grit value of the tactile standard while the unimodal visual matches were close to the grit value of the visual standard (i.e., steep VT exploration line in Figure 2C). This is in line with the notion that the experienced discrepancy was reduced in the oblique illumination condition because with decreasing experienced discrepancy when exploring the visuo-tactile standard bimodally, the need for compromise between the two senses also decreases. Interestingly, the results of Experiment 2 (Fig 2C) appear to more closely resemble the findings by Lederman and Abbott (see their Fig 2, page 907), which might indicate that our oblique illumination provided a closer approximation of participants’ visual experience in their Experiment 1.

Under both illumination conditions, participants quite accurately matched the visual and tactile standards in the unimodal exploration conditions to the correct matching stimulus within and across modalities. This seems, at first glance, to contradict the findings of Brown [21] and Ho et al. [23]. Brown [21] observed larger accuracy for visual performance under oblique illumination. While our task was not particularly suited to thoroughly investigate the accuracy of matching performance, our results reveal similar accuracy in matching the unimodal standards to the correct matching stimulus (i.e., 60 grit tactually and 150 grit visually) in the two illumination conditions. Furthermore, to concur with Ho et al., we might have expected the participants to match the visual standard to a rougher matching stimulus tactually (V-T condition), and the tactile standard to a less rough matching stimulus visually (T-V condition) under oblique illumination. Descriptively, the visual match was perceived as rougher after tactile exploration (T-V condition) in Experiment 2, but those observations were not statistically significant. However, in contrast to Ho et al., our comparisons were made cross-modally and, most importantly, based on a relatively coarse (and thus probably not sensitive enough) grit scale. The fact that the variability of weights (and of the matches) changed with illumination is, however, in line with the supposition that perceived roughness increased under oblique illumination and therefore indirectly corroborate the findings by Ho et al., that the perceived visual roughness changes with illumination angles.

Finally, we also wanted to explore if and how tactile weights correlate with haptic preferences and tactile acuity, especially in cases where we observed large differences in individual weights. For example, in Experiment 1, some participants seemed biased towards the visual or tactile modality to varying degrees. However, no correlations with tactile acuity measures and haptic preferences were observed. Together, with the lack of a correlation between the weights in the two experiments, this seems to speak against a consistent influence of individual modality preferences and acuity measures. This is also in line with previous studies that failed to establish a clear link between tactile acuity and tactile experience with the size of individual haptic biases [16, 22], as well as between tactile acuity and roughness discrimination performance [32]. Note though, that our sample size might have been too small to detect potential correlations, so it may be worth assessing this further in studies using larger samples.

## Conclusion

We replicated Lederman and Abbott’s [5] finding of an equal visuo-tactile contribution in surface texture perception on a group level under two different illumination conditions. Our within-subject design further revealed substantial inter-individual variations in modality weights. More specifically, we found that equal weighting of sensory information seems to hold only within a limited range of experienced discrepancy, and this range may vary between participants. Outside of this range, participants may no longer integrate information from different modalities, but may instead discount information from one modality (for a similar argument, see [12, 33]).

